# A Lamin Associated Chromatin Model for Chromosome Organization

**DOI:** 10.1101/788497

**Authors:** Ajoy Maji, Jahir Ahmed, Shubhankar Roy, Buddhapriya Chakrabarti, Mithun K. Mitra

## Abstract

We propose a simple model for chromatin organization based on the interaction of the chromatin fibres with Lamin proteins along the nuclear membrane. Lamin proteins are known to be a major factor that influences chromatin organization, and hence gene expression in the cells. Our polymer model explains the formation of lamin associated domains, and for heteropolymers with sequence control, can reproduce observed length distributions of LADs. In particular, lamin mediated interaction can enhance the formation of chromosome territories as well as the organization of chromatin into tightly packed heterochromatin and the loosely-packed gene-rich euchromatin regions.

## Introduction

The link between microscopic and macroscopic descriptions of genome structure and function is one of the key questions of present day biology [1–3]. In particular, the structure and folding principles of the interphase chromosomes has been the subject of much debate over the last decade[4–6]. While the structure of DNA double helix and histone proteins that form the nucleosome is well understood[7–9], how the nucleosomes finally organise to form the interphase chromosomes still remains an open question[10–12]. The organization of the chromatin determines the function of the cell type, with the epigenetic state of a differentiated cell correlated with differential packaging of the chromosome. Understanding the principle behind the chromatin organization thus has important implications for the proper functioning of the cell as misfolding errors leads to several human pathologies.

The advent of experimental techniques such as chromosome conformation capture (2C/3C/5C/Hi-C)[13–16] and FISH[17, 18] has provided a wealth of information on the large scale structure of the chromosome. A key experimental observation has been that different chromosomes segregate into different territories with minimal contact between them[19–21]. Additionally, the chromatin can also be classified into transcriptionally silent heterochromatin and gene expression active euchromatin, with heterochromatin regions being more tightly packed, and located preferentially in the nuclear periphery. The euchromatin, in contrast, is relatively loosely packed and located in the interior[22–25].

One of the key mediators in the organization of the genome are interactions between the nuclear lamina and chromatin. The nuclear lamina (NL) is a complex structure that acts as a scaffold for various proteins that regulate nuclear structure and function [26, 27]. Mammalian cells contain two B-type lamins, B1 and B2, and two A-type lamins, lamins A and C. In most cells, lamins B1 and B2 are concentrated along the NL in nculear periphery, while A-type lamins are also found in the nuclear interior[28, 29]. The lamin proteins in the NL play an important role in the organization of the chromosomes within the nucleus. Experimental studies have shown that certain regions of the chromosomes lie in close proximity to the lamina, and DamID experiments have been used to build a map of chromosome lamin interactions[26, 27, 30, 31]. These experiments reveal that there are large domains of chromosomal regions that have a high degree of affinity for nuclear lamins (called lamina-associated domains or LADs), alternating with regions of very low affinity. The LADs in the human genome can be very large, ranging from 0.1*Mb* − 10*Mb* in length[30, 31]. The LADs are associated with gene poor regions of the chromosome, with the mean gene density being around half of that in the non-LAD regions. Additionally, other gene activity markers, such as PolII and the histone mark H3K4me2 are also repressed within LADs, indicating that LADs represent a strongly repressive chromatin environment[26, 27, 30]. A large fraction of the human chromosome (≈ 40%) consists of LADs[30].

Subsequent experiments have also shown that the interaction between LADs and lamin proteins are stochastic in nature, with only a fraction of the total LADs being in contact with the NL in a given cell[31]. After the cell division process, a new subset of LADs can be in contact with the NL. ChIP-DamID experiments have shown that only those LADs which interact with the lamin proteins have enhanced levels of H3K9me2, which implies that this methylation mark status is also stochastic, and directly correlated with the LAD position[31]. These experiments conclusively show that LADs are positioned stochastically within the nucleus.

While there has been considerable theoretical progress in understanding the origins of the three-dimensional organization of chromosomes [32–37], an important missing ingredient in the proposed models has been the effect of the NL-chromatin interactions. In this letter we propose a simple polymer model that incorporates confinement as well as the attractive interactions between the chromosomes and the lamin proteins. The main results are as follows: (i) for a homopolymer we reproduce experimentally observed scaling of chromatin size as a function of base pairs and their associated contact probabilities, (ii) for a heteropolymer with sequence control we obtain length distributions of LADs, and (iii) for a mixture of homo and hetero-polymers we observe phase separation of chromosomes and formation of distinct territories. We thus demonstrate that a complete understanding of the folding principles of the chromosome need to incorporate this interaction for a cohesive picture.

## Model

We model the chromatin as a self-avoiding polymer chain of *N* beads, each of diameter *b*_0_, connected by harmonic springs. The polymer is confined within a sphere of radius *R*. The polymer consists of two types of beads, type *A* and type *B* (see Fig. 1). The inner surface of the sphere attracts beads of type A, mimicking lamin-chromatin interactions. The fraction of type A beads is denoted by *f*. The lamin interactions occur if a bead of type A lies within a cut-off distance *R*_*c*_ = *b*_0_ from the inner wall of the sphere. The energy of the polymer chain is thus given by the Hamiltonian,

**FIG. 1.**
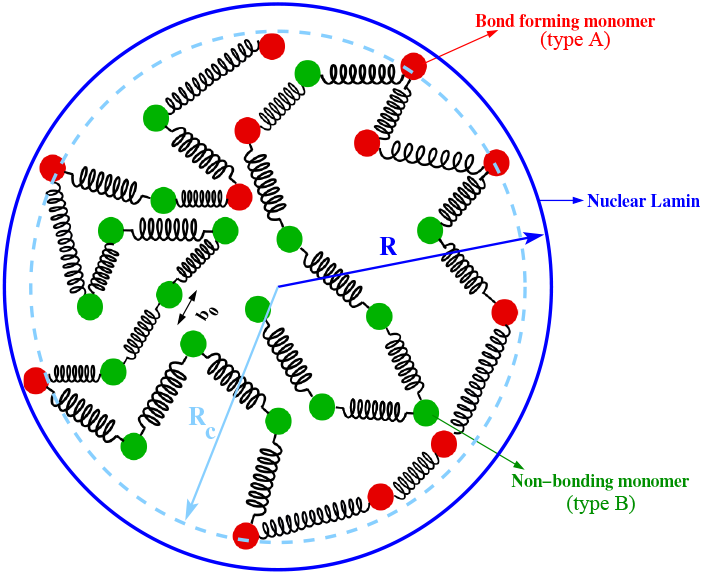
A schematic representation of a bead-spring polymer model with type A and type B beads confined within a sphere (see text)

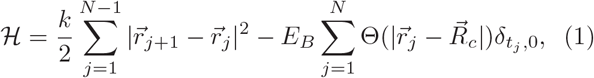

where *k* = 30 *k_B_T* denotes the spring constant of the bead-spring polymer, and is taken to be in the same range as previous studies [32, 37], and *t*_*j*_ denotes the type of bead (0 for A-type beads, and 1 for B-type beads). The lamin-chromatin binding energy is given by *E*_*B*_. Equilibrium conformations of this confined polymer system in the presence of an attractive boundary is then simulated following Metropolis Monte Carlo scheme, and the statistical averages are calculated for a time of *T*_*MC*_ = 10^9^ MC steps after allowing an equilibration time of *T*_*eq*_ ∼ 10*N* ^2^ ∼ 10^6^ MC steps.

The radius *R* of the nucleus and the polymer volume fraction is chosen in our model to conform to a biologically relevant scenario. Human nuclei sizes can range between diameters of 6*µm* − 11*µm*[38, 39], while the total number of base pairs in the human genome is of the order of 6 billion (6 × 10^9^*bp*)[38]. Assuming that a 30 *nm* chromatin bead has 3000*bp*[40, 41], the chromatin volume fraction of the human nuclei ranges from 0.004 to about 0.25. For our simulated confined polymer with *N* = 512 monomers, the radius of the confining sphere then corresponds to *R* ≃ 12 for a volume fraction of 0.004 and *R* = 7 for a volume fraction of 0.25. We look at physical quantities of interest for these two values of the radius *R* = 12 and *R* = 7 in this Letter.

## Results

We first consider a homopolymer with all type *A* beads that can bind to the lamin proteins, i.e. *f* = 1. This is done for both values of the nuclear radii used in our investigations, *R* = 7 and *R* = 12 for a range of binding energies 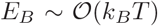. As shown in Fig. 2(a), the mean square displacement 〈*R*^2^(*s*)〉 as a function of contour length of the polymer *s* scales as 〈*R*^2^(*s*)〉 ∼ *s*^2*ν*^, with an exponent *ν* between 0.4 − 0.7 for short genomic distances and saturates for higher values. The saturation of the mean square separation is a simple consequence of the confinement of the polymer. For *R* = 12, *ν* ≈ 0.5−0.7; while for *R* = 7, *ν* ≈ 0.4−0.6 which corresponds extremely well with measured values from FISH data[42–44].

**FIG. 2.**
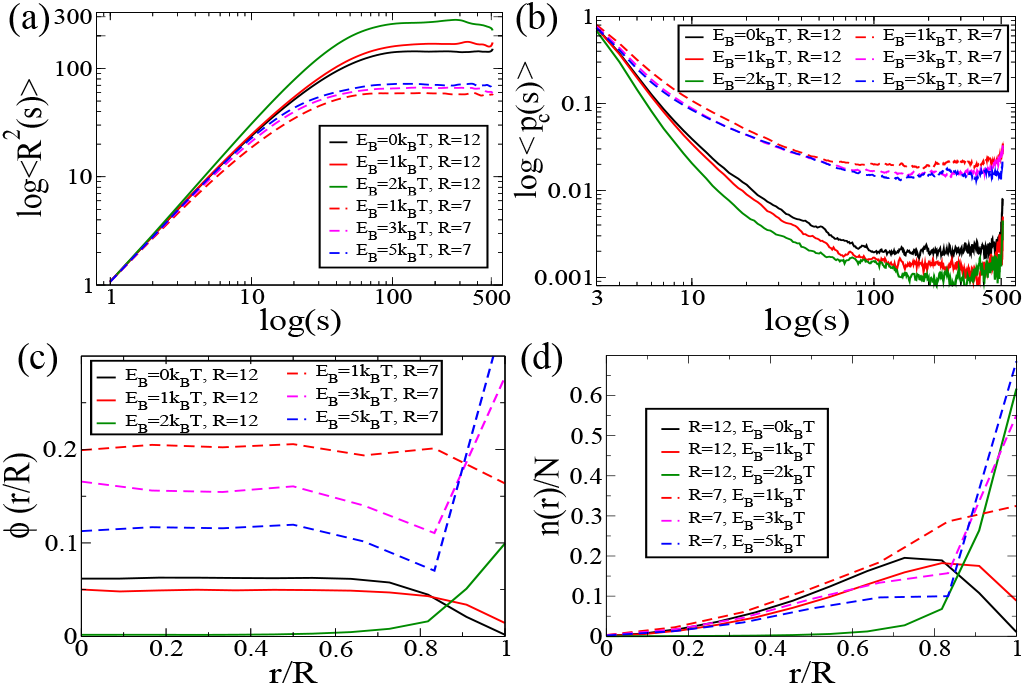
(a) Mean-square-distance 〈*R*^2^(*s*)〉 as a function of the separation *s* for different binding energies and different volume fractions; (b) Contact probability *p*_*c*_(*s*) as a function of the separation *s* for different binding energies and different volume fractions; (c) The volume fraction within spherical shells as a function of the normalised radial distance; (d) The (normalised) number of monomers within a spherical shell as a function of the normalised radial distance

We also compute the contact probability for two monomers separated by *s* base pairs to approach within a certain cutoff distance *r*_*cut*_ of each other. Consistent with previous studies, we choose *r*_*cut*_ = 2.5*b*_0_ [34]. For small base pair separations, the contact probability decreases before tapering off at large values (see Fig. 2(b)). The contact probability decreases following a power law of the form *p*_*c*_(*s*) = *s*^−*β*1^ for small values of *s*. The exponent *β*_1_ ≈ 1.5 − 1.6 for *R* = 12 and ≈ 0.9 − 1.0 for *R* = 7, consistent with Hi-C experiments[45, 46].

Further, we compute volume fraction of monomers as a function of the radial distance from the centre, i.e. we count the number of monomers *n*_*r*_ in a thin shell *r* → *r* + ∆*r*, with ∆*r* = 1, and compute the volume occupied by these *n*_*r*_ monomers normalized by the volume of the shell. This is shown in Fig. 2c. The corresponding number of monomers (normalised by the total number *N*) in each shell is shown in Fig. 2(d). Entropic confinement, in the absence of lamin-protein interactions (*i.e. E*_*B*_ = 0), leads to a low volume fraction near the surface and a constant value throughout the rest of the nucleus. For *E*_*B*_ ≥ 2 *k_B_T*, the chromatin volume fraction at the nuclear periphery increases such that the outermost shell is more densely packed than the inner ones. This is consistent with observations of tighter chromatin packing in heterochromatin regions[22–25], and also correlates with the hypothesis that lamin protein chromatin interactions are more prominent in heterochromatin regions, which explains their tighter packing, and consequently, higher volume fractions.

Next, we investigate domain formation within our model to compare against DamID experimental data. We define the lamin proximity index for the *i*^*th*^ monomer as, 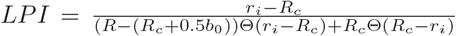, which determines the position of the centre of a monomer, with respect to the distance *R*_*c*_ beyond which it interacts with the lamin. The *LP I* value ranges from [−1: 1], with positive values indicating bond-formation with the lamin. Figures 3(a,b) and (d,e) show the variation of *LP I* as a function of the binding energy *E*_*B*_ for both spheres of radii *R* = 12 and *R* = 7 respectively.

**FIG. 3.**
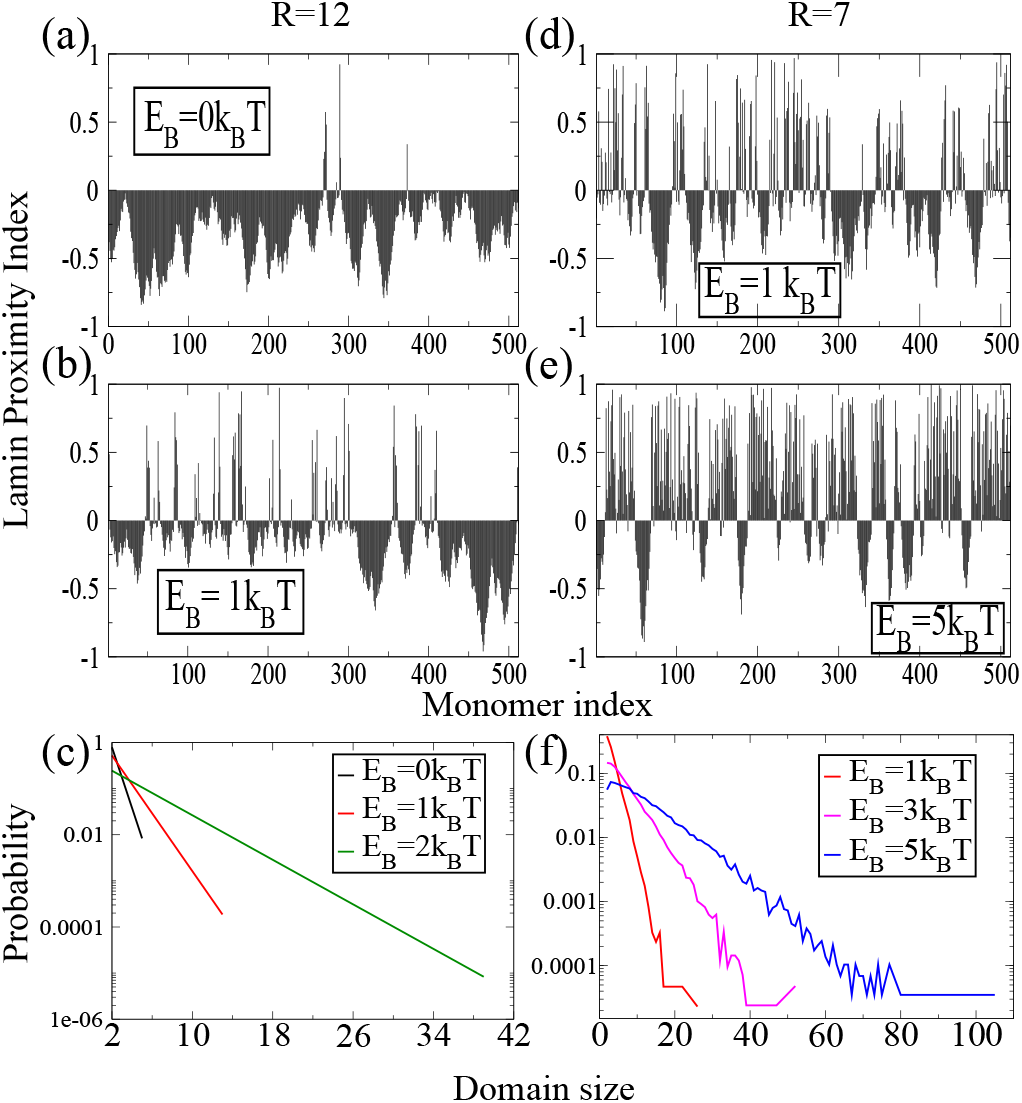
Lamin proximity indices for R=12 and binding energies (a) *E*_*B*_*T* = 0*k*_*B*_*T* and (b) *E*_*B*_ = 1*k*_*B*_*T*; LPI for R=7 and binding energies (d) *E*_*B*_ = 1*k*_*B*_*T* and (e) *E*_*B*_ = 5*k*_*B*_*T*; (c)&(f): Domain size distributions for different binding energies for R=12 and R=7 respectively

The lamin-associated chromatin domains for the homopolymer, has a size distribution *P* (*ℓ*) ∼ exp[−*ℓ/ℓ*_0_], peaked around *ℓ* = 0, with the characteristic length scale *ℓ*_0_ monotonically increasing as a function of the binding energy *E*_*B*_ (Fig. 3(c,f)). The size-distribution obtained from DamID data is peaked around a non-zero value of *ℓ* [30]. This arises due to the fact that for chromatin roughly 40% of the monomers can associate with the lamin. This necessitates a heteropolymer model of chromatin where only a fraction *f* of the beads (type *A*) can associate with the lamin proteins.

## Heteropolymer

A heteropolymer model for chromatin assumes *N*_*A*_ beads of type *A* and *N*_*B*_ beads of type *B*, such that *N_A_/*(*N*_*A*_ + *N*_*B*_) = *f*. We consider three heteropolymer models: *(i)* random-heteropolymer - where the *N*_*A*_ number of *A* types beads are chosen randomly; (ii) uniform-block-copolymer - where the *N*_*A*_ beads are divided into uniform patches of size *p* = *N/*4; and *(iii)* gaussian-block-copolymer - where the *N*_*A*_ beads are divided into patches with patch sizes are chosen from a Gaussian distribution (*µ* = 20*, σ* = 5). For polymers with quenched randomness of the type studied here, the fraction *f* as well as the disorder-correlation length plays a role in dictating their equilibrium properties. We investigate the statistical properties of the confined polymer both as a function of the lamin binding energy as well the fraction *f* of binding monomers.

The mean square displacement between any two monomers 〈*R*^2^(*s*)〉 has the same statistical features as a homopolymer, with 〈*R*^2^(*s*)〉 ∼ *s*^*ν*^ for small separations followed by a saturation at larger values. The exponent *ν* increases as a function of *E*_*B*_ for all fractions considered, as shown in Fig. 4(a). Increasing the binding energy *E*_*B*_ allows the monomers to spread out on the surface, as lamin attachments become more favourable, leading to an increase in the exponent *ν*. As expected, *ν* also increases with increasing the fraction *f* of binding monomers. A similar behaviour is observed for the contact probability exponent *β* (Fig. 4b). The values of the exponents *ν* and *β* are similar for the different disorder realisations studied.

**FIG. 4.**
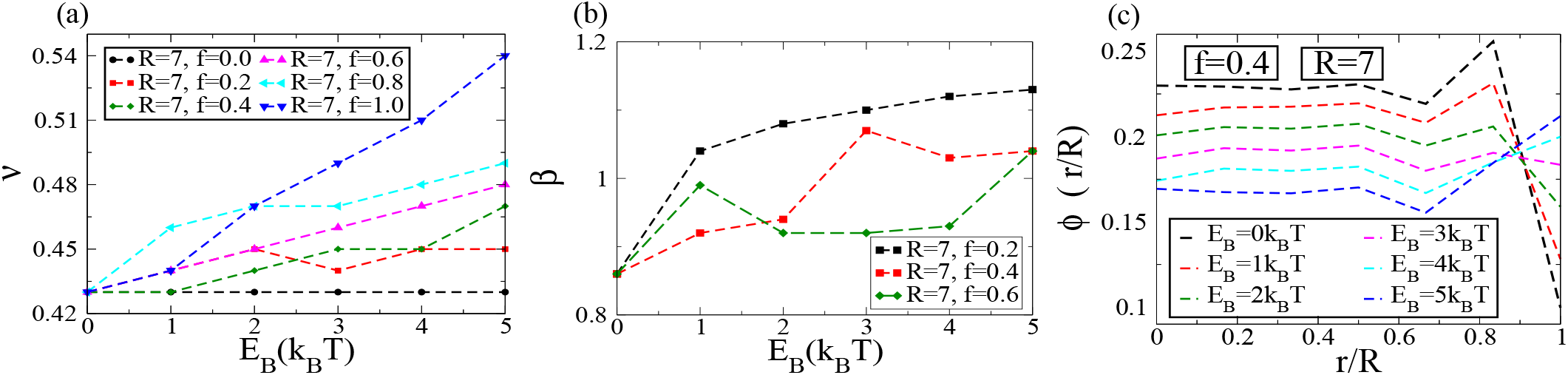
(a)The mean square displacement exponent *ν* as a function of *E*_*B*_ for different fraction of binding monomers *f*. (b) The contact probability exponent *β* as a function of *E*_*B*_ for different *f*. (c) Plots of the radial volume fraction. All results are shown for the random heteropolymer model in a spherical cavity of *R* = 7

We also plot the volume fraction (Fig. 4c) as a function of the normalised distance from the center of the sphere, for a binding fraction of *f* = 0.4 corresponding to the biologically relevant situation. For small binding energies, the volume fraction drops close to the surface of the nucleus. With increasing binding energy the volume fraction shows a maxima near the lamina, indicating an increased density of monomers there.

We now turn to the lamin contact maps in order to investigate domain formation for the heteropolymer case. A representative plot for the Lamin Proximity Index (*LP I*) for a *random* heteropolymer with *f* = 0.4 is shown in Fig. 5(a) for *R* = 7 and *E*_*B*_ = 3*k_B_T*. Domains of lamin-associated-chromatin alternate with ones that do not come in contact with the NL. The length distribution of the domains *P* (*ℓ*) for *R* = 7 and *R* = 12 are shown in Figures 5(b) and 5(c) respectively. The domain sizes are distributed exponentially for both values of the nuclear size indicating that a random heteropolymer model does not reproduce the characteristic distributions of LADs observed in experiments [30]. Fig. 5d) shows the *LP I* for the uniform block copolymer with *f* = 0.5 having 4 equal length alternating blocks of attractive and inert domains that interact with the lamin. The block copolymeric nature is reflected in the LPI plots (Fig. 5d) with larger domain sizes in comparison to those formed in the case of the random heteropolymer. The associated domain length distributions for *R* = 7 and *R* = 12 are shown in Figs. 5 (e) and (f) respectively. In contrast to the single exponential fit for both the homopolymer and the random heteropolymer, the domain size distribution in this case is a double exponential, with an enhanced probability for larger domains. For high enough binding energies, additionally there is a peak corresponding to the domain size as well. Finally we consider the case of Gaussian block heteropolymer with *f* = 0.4, with the corresponding LPI shown in Fig. 5(g). The distributions of the LAD sizes reflect the Gaussian nature of the block copolymer for higher values of the binding energy *E*_*B*_ ≳ 2*k_B_T*, as can be seen from Figs. 5(h) and (i) for both values of the radius. This qualitatively agrees with the domain size distribution observed in DamID measurements[30]. The heteropolymer model thus illustrates that the observed domain length distributions in experimental studies must necessarily correlate with the distribution of chromatin regions that can interact with lamin proteins as well as setting a scale for the strength of the lamin-chromatin interactions. Weak interactions give rise to exponential domain length distributions, and hence the interaction energies must be beyond a certain strength in order to explain the experimental observations.

**FIG. 5.**
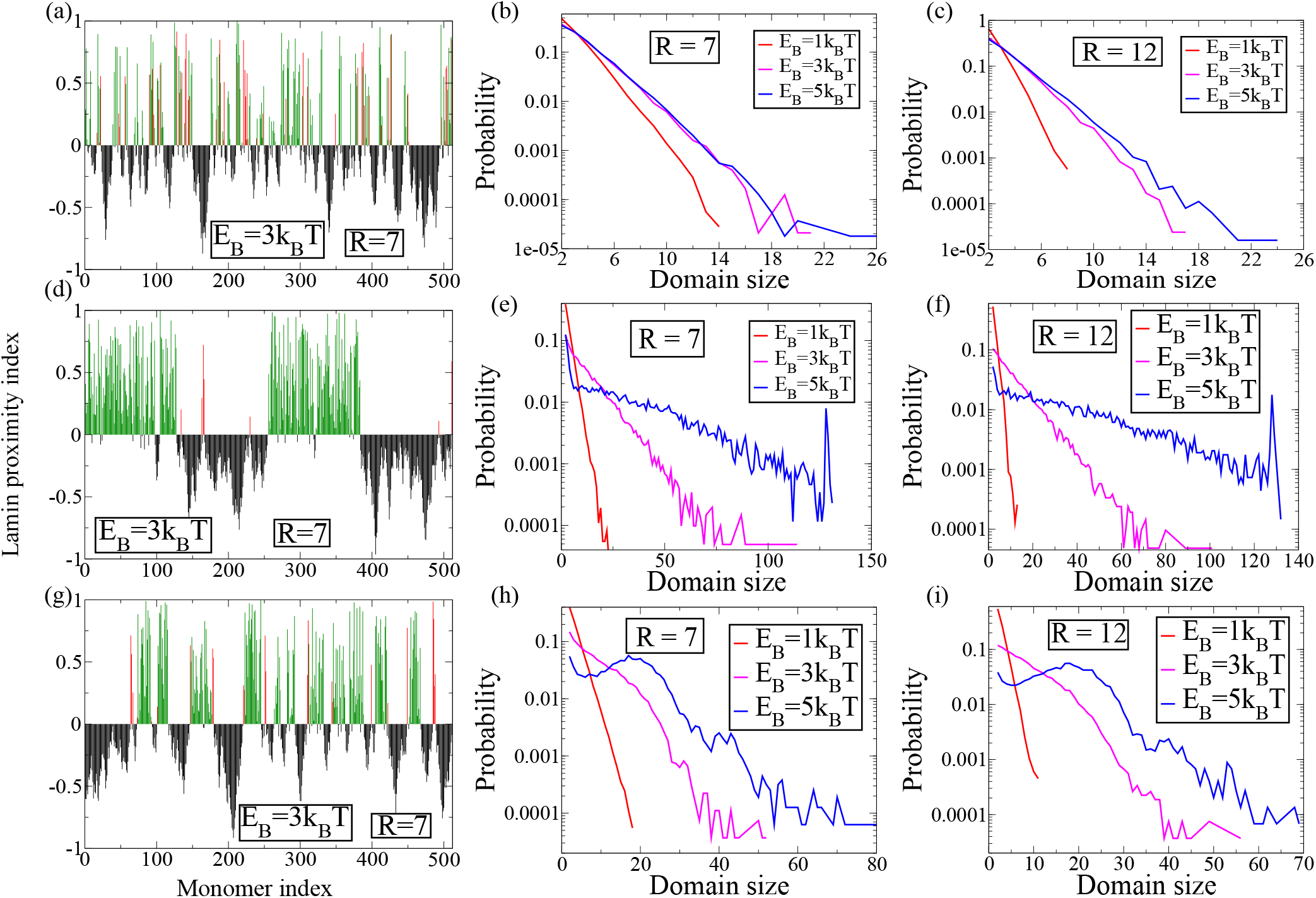
Lamin proximity index for *E*_*B*_ = 3*k*_*B*_*T* for a sphere with *R* = 7 for the three heteropolymer models: (a) random heteropolymer, (d) uniform block copolymer, and (g) Gaussian block copolymer. The red lines indicate type *B* monomers that are in proximity to the NL, while green lines indicate bond-forming type-A monomers. The corresponding distributions of domain sizes is shown for *R* = 7 in panels (b, e and h), and for *R* = 12 in panels (c, f and i).

## Multiple polymers

We explore the formation of chromatin territories within our model. In the absence of any lamin interactions (*E*_*B*_ = 0), the lamin contact maps (Fig. 6a) shows negligible territory formation. A sample equilibrium configuration, shown in Fig. 6(c) illustrates that there is significant interpenetration between the four polymer strands. If we allow two of the four polymers to interact with the lamin, these two polymers show extremely low levels of interpenetration. This can be seen from the contact map shown in Fig. 6(b), where the first polymer (monomers 0 − 127), and the third polymer, (monomers 256 − 383) interacts with the lamin (*E*_*B*_ ≠ 0), while the other two do not. The regions of the contact map corresponding to overlaps between the first and third polymers clearly show a reduced intensity, indicating the two polymers seldom approach each other. This is also clear in the sample equilibrium configuration shown in Fig. 6(d), where the first (brown) and third (red) polymers stay close to the attractive surface on the two distal sides of the sphere. The two noninteracting polymers (shown here in cyan and blue) on the other hand, occupy the central region of the sphere and have increased inter-penetration. The interactions of the lamin proteins with the chromatin thus enhances the ability of the chromatin to segregate into individual territories. The attractive surface interaction makes it energetically favourable for the chromatin polymers to occupy different regions of the nucleus, leading to the formation of territories. This provides a candidate mechanism by which chromosome territories form within the eukaryotic cell nucleus.

**FIG. 6.**
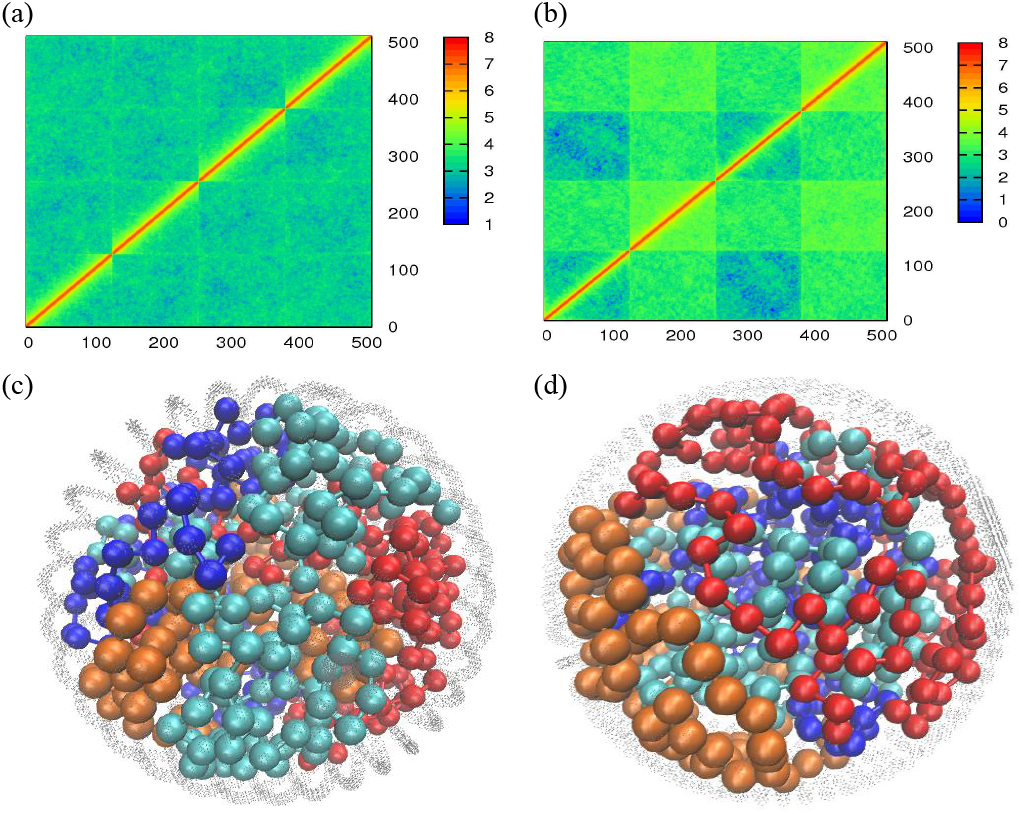
(a) Contact maps and (c) sample configuration for system of 4 polymers, none of which interact with lamin proteins. (b) Contact maps and (d) sample configuration for a system of 4 polymers of which two interact with lamin proteins while the other two do not.

## Discussion

In conclusion, we model chromatin packing in the cell nucleus as a polymer confined in a spherical cavity having an attractive interaction with the inner walls. We compute statistically measurable quantities such as mean square displacement as a function of base pair separation 〈*R*^2^(*s*)〉, the lamin proximity index *LP I*, and contact map for homo and heteropolymers with different disorder realisations and different nuclear radii. Our computational results explain the data observed in DamID and FISH experiments.

A complete map of chromatin-nuclear lamina interactions generated by DamID experiments have identified a large number of LADs in the human chromosome, and understanding the origin of the observed distribution of LADs is thus important to understand the 3D organization of the genome. Our work shows that a heteropolymeric chromatin, with domains of lamin-binding regions with sizes drawn from a distribution, reflects this structural heterogeneity. We show, for example, that for Gaussian distributed domains, the lengths of lamin associated regions also follow a Gaussian distribution. Further, for low binding energies, the distribution of lengths follows a single exponential. Thus our analysis shows that lamin chromatin interactions need to be beyond a certain critical strength (few *k_B_T* s), such that thermal energies can compete with the interaction energy leading to dynamic chromosomes, and also reflect the underlying domain structure.

For the case of multiple polymers, while it is known that confinement can introduce territory formation [47], we show that lamin mediated interactions can effectively strengthen the formation of chromosome territories, even at densities where territory formation would not be expected from simple confinement.

While our model shows several interesting features a quantitative understanding of genome organization is far from complete. We assumed an uniformly attractive surface, corresponding to a uniform distribution of lamin proteins. In reality, there is a complex and heterogeneous distribution of lamin filaments on the nuclear envelope, and this would lead to tighter heterochromatin packaging at regions of high lamin concentration [48]. Further, A-type lamins are known to be present on the nucleolas as well[28, 29]. Thus LADs on chromatin can interact with the nucleolas surface in the interior of the nucleus. The effect of this attractive surface in the nuclear interior can compete with the attractive interaction at the nuclear periphery to give rise to non-trivial structures. The current model focuses solely on the effect of nuclear lamin interactions. However, it is known that various other proteins such CTCF[49, 50], cohesins and condensins[51–53] play a role in the large-scale organization of the genome. An open question is how protein-mediated interactions in the bulk compete with lamin-interactions on the surface to determine 3D organization. Non-equilibrium active forces may also play a role, although their role in large scale organization remains open. While preparing this manuscript for publication we became aware of a similar model by Chiang *et al.*[54]. While our model does not include sequence specific data as an input into hetero/euchromatin track lengths, it is more generic, allowing for systematically tuning the disorder correlation length of heterochromatin tracks along the backbone. Despite its conceptual simplicity, our model captures features of conformational statistics of the chromatin. This calls for a systematic study of specific interactions among beads and lamin to understand robustness of genetic landscape on changes in mesoscale parameters.

We hope that our simple theoretical model will encourage experimental work in this direction. Our study predicts that tuning the strength of lamin-chromatin interactions can change the distributions of lamin-associated domain lengths. If the interactions can be made sufficiently weak, the theory predicts a transition to a single exponential distribution of domain sizes. Further, tuning the interaction strength will also change the radial volume fraction, with volume fraction at the nuclear periphery increasing monotonically with increasing interaction strength. These specific predictions can be tested against experimental data.

Our study emphasizes how sequence heterogeneity in the genome affects the three-dimensional genome organization. In particular, this heterogeneity is crucial to a complete understanding of the chromosome packaging problem.

## Acknowledgements

Financial support is acknowledged by MKM for Ramanujan Fellowship (13DST052), DST and IITB (14IRCCSG009).

## References

[1] P. Fraser and W. Bickmore, Nature 447, 413 (2007).

[2] T. Misteli, Cell 128, 787 (2007).

[3] C. Lanctot, T. Cheutin, M. Cremer, G. Cavalli, and T. Cremer, Nature Reviews Genetics 8, 104 (2007).

[4] R. A. Horowitz-Scherer and C. L. Woodcock, Chromosoma 115, 1 (2006).

[5] K. Bystricky, P. Heun, L. Gehlen, J. Langowski, and S. M. Gasser, Proceedings of the National Academy of Sciences 101, 16495 (2004).

[6] T. Cremer and C. Cremer, Nature reviews genetics 2, 292 (2001).

[7] H. Schiessel, W. M. Gelbart, and R. Bruinsma, Biophysical Journal 80, 1940 (2001).

[8] A. Annunziato, Nature Education 1, 26 (2008).

[9] L. Mariño-Ramïrez, M. G. Kann, B. A. Shoemaker, and D. Landsman, Expert review of proteomics 2, 719 (2005).

[10] K. Van Holde, J. Allen, K. Tatchell, W. Weischet, and D. Lohr, Biophysical journal 32, 271 (1980).

[11] J. Finch and A. Klug, Proceedings of the National Academy of Sciences 73, 1897 (1976).

[12] C. Woodcock, L.-L. Frado, and J. Rattner, The Journal of cell biology 99, 42 (1984).

[13] J. Dekker, K. Rippe, M. Dekker, and N. Kleckner, science 295, 1306 (2002).

[14] J. Dekker, Nature methods 3, 17 (2006).

[15] O. Oluwadare, M. Highsmith, and J. Cheng, Biological procedures online 21, 7 (2019).

[16] J. Han, Z. Zhang, and K. Wang, Molecular Cytogenetics 11, 21 (2018).

[17] J. M. Levsky and R. H. Singer, Journal of cell science 116, 2833 (2003).

[18] L. Giorgetti and E. Heard, Genome biology 17, 215 (2016).

[19] T. Cremer and M. Cremer, Cold Spring Harbor perspectives in biology 2, a003889 (2010).

[20] K. J. Meaburn and T. Misteli, Nature 445, 379 (2007).

[21] W. A. Bickmore, Annual review of genomics and human genetics 14, 67 (2013).

[22] S. T. Kosak and M. Groudine, Genes & development 18, 1371 (2004).

[23] P. Meister, B. D. Towbin, B. L. Pike, A. Ponti, and S. M. Gasser, Genes & development 24, 766 (2010).

[24] P. K. Geyer, M. W. Vitalini, and L. L. Wallrath, Current opinion in cell biology 23, 354 (2011).

[25] E. Fedorova and D. Zink, Biochimica et Biophysica Acta (BBA)-Molecular Cell Research 1783, 2174 (2008).

[26] B. Van Steensel and A. S. Belmont, Cell 169, 780 (2017).

[27] I. Solovei, K. Thanisch, and Y. Feodorova, Current opinion in cell biology 40, 47 (2016).

[28] T. Dechat, A. Gajewski, B. Korbei, D. Gerlich, N. Daigle, T. Haraguchi, K. Furukawa, J. Ellenberg, and R. Foisner, Journal of cell science 117, 6117 (2004).

[29] R. D. Moir, M. Yoon, S. Khuon, and R. D. Goldman, The Journal of cell biology 151, 1155 (2000).

[30] L. Guelen, L. Pagie, E. Brasset, W. Meuleman, M. B. Faza, W. Talhout, B. H. Eussen, A. de Klein, L. Wessels, W. de Laat, et al., Nature 453, 948 (2008).

[31] J. Kind and B. van Steensel, Nucleus 5, 124 (2014).

[32] J. Mateos-Langerak, M. Bohn, W. de Leeuw, O. Giromus, E. M. Manders, P. J. Verschure, M. H. Indemans, H. J. Gierman, D. W. Heermann, R. Van Driel, et al., Proceedings of the National Academy of Sciences 106, 3812 (2009).

[33] L. A. Mirny, Chromosome research 19, 37 (2011).

[34] M. Barbieri, M. Chotalia, J. Fraser, L.-M. Lavitas, J. Dostie, A. Pombo, and M. Nicodemi, Proceedings of the National Academy of Sciences 109, 16173 (2012).

[35] N. Ganai, S. Sengupta, and G. I. Menon, Nucleic acids research 42, 4145 (2014).

[36] G. Gürsoy, Y. Xu, A. L. Kenter, and J. Liang, Nucleic acids research 42, 8223 (2014).

[37] H. Kang, Y.-G. Yoon, D. Thirumalai, and C. Hyeon, Physical review letters 115, 198102 (2015).

[38] B. Alberts, A. Johnson, J. Lewis, M. Raff, K. Roberts, and P. Walter, Molecular Biology of the Cell (Textbook). 5th ed. New York: Garland Science, 1115 (2008).

[39] H. B. Sun, J. Shen, and H. Yokota, Biophysical journal 79, 184 (2000).

[40] G. Wedemann and J. Langowski, Biophysical journal 82, 2847 (2002).

[41] S. Gerchman and V. Ramakrishnan, Proceedings of the National Academy of Sciences 84, 7802 (1987).

[42] S. Jhunjhunwala, M. C. van Zelm, M. M. Peak, S. Cutchin, R. Riblet, J. J. van Dongen, F. G. Grosveld, T. A. Knoch, and C. Murre, Cell 133, 265 (2008).

[43] L. S. Shopland, C. R. Lynch, K. A. Peterson, K. Thornton, N. Kepper, J. von Hase, S. Stein, S. Vincent, K. R. Molloy, G. Kreth, et al., The Journal of cell biology 174, 27 (2006).

[44] C. Münkel, R. Eils, S. Dietzel, D. Zink, C. Mehring, G. Wedemann, T. Cremer, and J. Langowski, Journal of molecular biology 285, 1053 (1999).

[45] F. Bantignies and G. Cavalli, Trends in Genetics 27, 454 (2011).

[46] R. Kalhor, H. Tjong, N. Jayathilaka, F. Alber, and L. Chen, Nature biotechnology 30, 90 (2012).

[47] S. Jun, A. Arnold, and B.-Y. Ha, Physical review letters 98, 128303 (2007).

[48] A. S. Belmont, Y. Zhai, and A. Thilenius, The Journal of cell biology 123, 1671 (1993).

[49] J. R. Dixon, S. Selvaraj, F. Yue, A. Kim, Y. Li, Y. Shen, M. Hu, J. S. Liu, and B. Ren, Nature 485, 376 (2012).

[50] S. S. Rao, M. H. Huntley, N. C. Durand, E. K. Stamenova, I. D. Bochkov, J. T. Robinson, A. L. Sanborn, I. Machol, A. D. Omer, E. S. Lander, et al., Cell 159, 1665 (2014).

[51] J. Lee, Journal of Reproduction and Development 59, 431 (2013).

[52] K. C. Yuen and J. L. Gerton, PLoS genetics 14, e1007118 (2018).

[53] F. Uhlmann, Nature reviews Molecular cell biology 17, 399 (2016).

[54] M. Chiang, D. Michieletto, C. A. Brackley, N. Rattanavirotkul, H. Mohammed, D. Marenduzzo, and T. Chandra, Cell reports 28, 3212 (2019).

